# A rapid approach for linear epitope vaccine profiling reveals unexpected epitope tag immunogenicity

**DOI:** 10.1101/2024.12.08.627427

**Authors:** Kirsten Browne-Cole, Kyrin R. Hanning, Kevin Beijerling, Meghan Rousseau, Jacelyn Loh, William Kelton

## Abstract

Antibody epitope profiling is essential for assessing the robustness of vaccine-induced immune responses, particularly while in development. Despite advancements in computational tools, high throughput experimental epitope validation remains an important step. Here, we describe a readily accessible method for rapid linear epitope profiling using phage-displayed oligo pools in combination with Nanopore deep sequencing. We applied this approach to TeeVax3, a Group A Streptococcus vaccine candidate, to investigate the antibody response generated in a pre-clinical rabbit model and assess antigen immunogenicity. Surprisingly, we found a strong bias in antibody binding response towards the N-terminal epitope tag used for purification. These tags are widely reported to have low immunogenicity and are frequently left uncleaved in pre-clinical studies. We further confirmed that the observed immune response against the epitope tag dominated even the conformational binding response and, using synthetic peptides, narrowed the epitope down to a set of 10 residues inclusive of the Histidine residues. Our findings highlight the importance of epitope-tag removal in pre-clinical studies and demonstrate the utility of rapid nanopore sequencing for early-stage vaccine evaluation.

## Introduction

Vaccine protection is strongly correlated with the quality and longevity of B and T cell responses induced against a given pathogen^1–3^. While T cell responses are driven by MHC loading of linear peptide epitopes present in vaccine antigens, antibodies can engage targets directly by binding to either linear epitopes or conformational epitopes formed when antigenic moieties are brought into close proximity by three-dimensional folding. Understanding the epitope distribution and binding affinity of the antibody response provides invaluable information for vaccine development efforts, especially where responses towards specific neutralizing epitopes are desired^4,5^. Alongside predictive computational tools^6^, several high-throughput wet lab approaches are widely used to profile the antigenic epitopes driving antibody responses^7,8^, particularly for applications in virology. These approaches can be adapted more widely to vaccine profiling^9^.

Crystallography remains the highest accuracy technique for antibody epitope mapping, followed by more recently developed Cryo-EM approaches^10^. Unfortunately, throughput is a challenge for these methods, and capturing a comprehensive snapshot of an antibody response against an antigen is very resource intensive. Instead, protein display systems have gained popularity as millions of individual epitope sequences can be physically linked to an underlying genotype that can be recovered by next-generation sequencing. Antigens can be expressed as full-length proteins or tiled as short peptides. The VirScan approach used phage display to present more than 1 x 10^8^ unique linear peptides from human viruses for capture by sera-derived antibodies from a global cohort of individuals^11,12^. In addition to antigen tiling, peptide sequences can also be systematically mutated using techniques like deep mutational scanning to provide greater precision when defining antibody epitopes^13,14^. The use of next-generation sequencing is particularly advantageous as it allows the detection of rare binding events at low frequencies, important when defining the binding ranges of polyclonal antibodies.

Here, we used a peptide phage display approach to investigate antibody responses elicited by TeeVax3^15^, a component of the multivalent vaccine candidate under development for Group A Streptococcus (Strep A) infection. Despite longstanding development campaigns by academic and industry groups^16^, no vaccines have yet been approved for this pathogen, in part because of large strain diversity and the risk of autoimmune induction by surface antigens (e.g. M protein). By instead concatenating sets of five to seven T antigens from diverse strains into three recombinant proteins (e.g. TeeVax3: T6M-T2M-T25M-T23M-T4N), protective immunity was induced in small animal models of Strep A infection^15^. Further work characterizing the antibody response in mice and rabbits used phage-displayed Fab immune libraries to isolate and crystallize high-affinity Fab domains against the T antigen^17^. High-resolution epitope mapping of antibody repertoires is key to understanding the drivers of vaccine efficacy. Surprisingly, there appeared to be immunodominant conformational epitopes towards the N-terminus of some T antigens with unusual temperature-dependent accessibility at 37 °C.

To better understand antibody biases and binding distribution on the TeeVax3 antigen in high throughput, we created a bacteriophage-based method for the rapid profiling of linear epitopes using Nanopore sequencing. Applying our method to sera from TeeVax3 immunized rabbits revealed a strong bias towards the N-terminal TEV/6-His tag sequence that dominated even the conformation antibody binding response. While epitope tags are routinely removed from vaccines, including most but not all TeeVax3 animal experiments^15,17^, conventional wisdom suggests these peptides should possess low immunogenicity^18,19^. In many cases, these tags are left on vaccine antigens, especially through the early stages of *in vivo* profiling in animal models^20–22^. As well as providing an updated set of tools for linear epitope profiling, our finding provides a cautionary tale for those developing antigen-based vaccines requiring exogenous peptide tags for purification or antigen tracking.

## Results

### Construction of tiled TeeVax peptide libraries for phage display

To profile TeeVax3 linear epitopes in high throughput, we developed a phage-based peptide display approach to make use of rapid Nanopore sequencing (**Figure 1a**). This sequencing approach can have higher error rates than other next-generation sequencing approaches, such as Illumina, but we reasoned that the unique offset nature of the peptide tiles enables identification regardless of sequencing error. We first cloned short control peptides encoding the 6-His tag and a cyclic peptide mimetic of lipooligosaccharide derived from the pathogen *N. gonorrhea* (PEP-1)^23^ into pAK200 phage display vectors^24^. These peptides were selected as high-affinity antibodies targeting them are readily available. ELISA analysis confirmed control phage binding to both anti-His and PEP-1 (mAb 2C7^25^) monoclonal antibodies above background, despite some non-specific binding being detected for the anti-His clone (**Figure 1b**). Having demonstrated the suitability of this system for the accessible display of short peptides, we next created a library of peptides from the 940 amino acid TeeVax3 antigen. Bioinformatic tiling was used to generate 308 sequences, each 20 amino acids in length and offset by three amino acids, except for a final tile with a two amino acid offset. These sequences were back translated, adapter sequences added to the 5’ and 3’ ends, and the resulting library ordered as an oligo pool (Twist Biosciences). A single cycle of PCR was used to create the second strand of DNA for restriction enzyme cloning into pAK200 and phage production.

**Figure 1:**
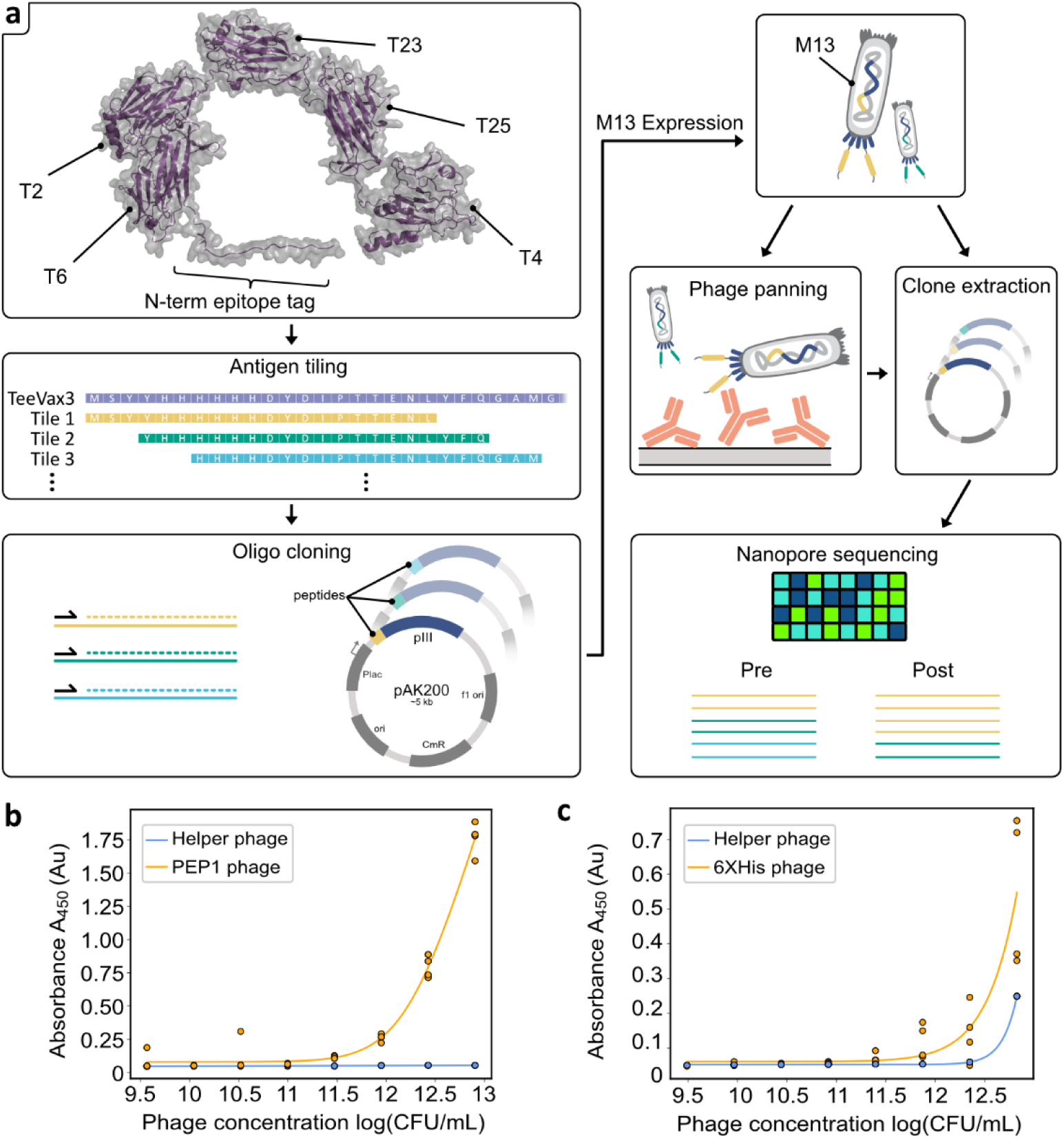
Linear antibody epitope profiling using phage panning and Nanopore sequencing. **(a)** Overall experimental scheme. Target antigen sequences are tiled into discrete peptides and encoded as an oligonucleotide pool. After a single cycle of PCR to create the complementary strands, these sequences are cloned into pAK200 plasmids and expressed on the pIII domain of M13K07 bacteriophage. Peptide tile abundances before and after panning against the vaccine-specific antibody pool are determined by Nanopore sequencing and the ratio of these frequencies used to determine epitope enrichment. ELISA analysis was used to show control phage expressing model PEP1 cyclic peptides **(b)** and 6-His epitope tag peptide **(c)** bind to 2C7 and anti-His monoclonal antibodies respectively (n = 4) relative to helper phage without peptide expression.

### Characterization of TeeVax3 peptide libraries by Nanopore sequencing

Prior to phage epitope profiling, we first confirmed the cloned library coverage using Nanopore sequencing. Plasmid DNA was isolated and cloned amplicons were removed by restriction enzyme digest to eliminate PCR amplification bias. We obtained ≥1.4 million reads from the library using a Q score cutoff of 15 (**Supplementary Fig. S1a)** with a median fragment length of 720 bp corresponding to the expected size for the digested fragment (**Supplementary Fig. S1b**). Reads were then binned into tiles via Minimap2. We again observed that 100% of the expected tiles were present, although individual tile frequencies within this pool ranged from 0.07% to 0.57% (Figure 2a). Bacteriophage were then produced and immediately used to reinfect cells to gather the fraction of the library amenable to expression by phage. Clonal dropout at this step would lead to gaps in epitope coverage; therefore, nanopore sequencing was again used to re-evaluate tile frequencies. We again sequenced to a depth of ≥1.4 million reads and saw 100% retention of tile coverage in the phage library. Here, we observed minor differences in tile frequency and a general broadening of the tile abundance distribution when compared to Round 0 (Pre-expression plasmid sequence pool). Individual tile frequencies ranged from 0.036% to 1.82% indicating some expression biases but this was not concentrated to any one region of the antigen (Figure 2b).

**Figure 2:**
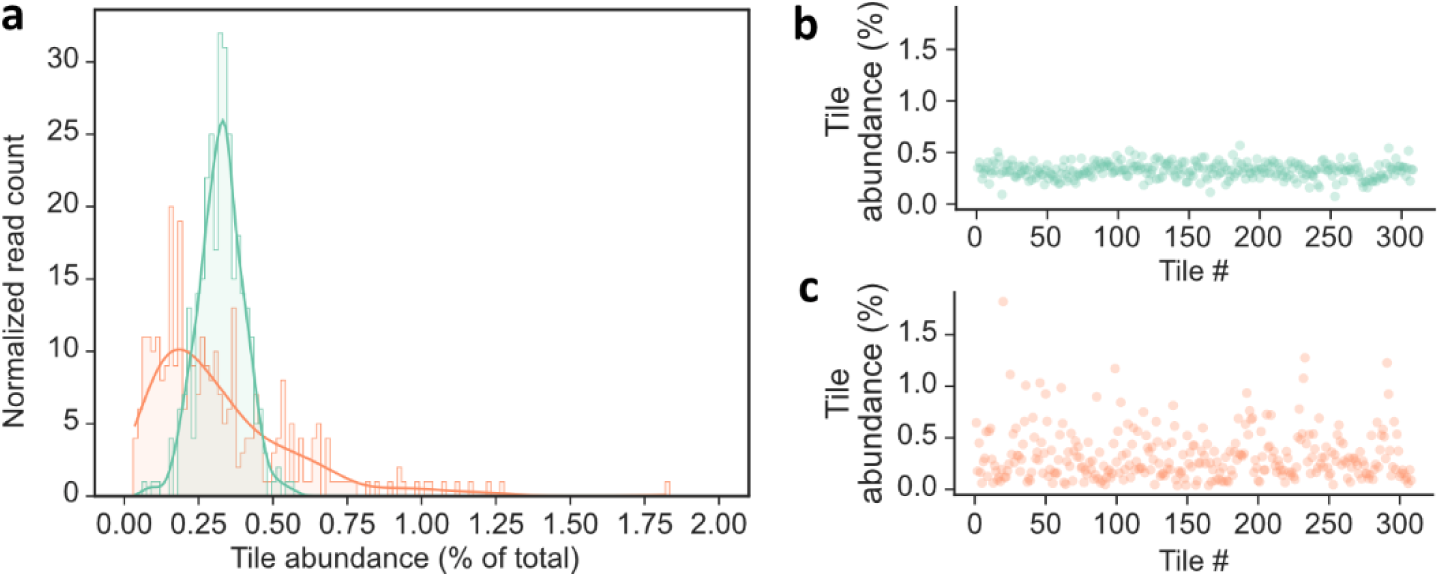
Pre-panning peptide library evaluation by Nanopore Sequencing. **(a)** Distribution of individual tile abundances before (green) and after (orange) expression on phage, normalized to the total read count. Solid lines represent smoothed distributions generated by kernel density estimations of the underlying histogram plots. **(b)** To determine whether expression bias is distributed evenly throughout the antigen, individual tile abundances were plotted against tile location within the antigen (Tile #) for Round 0 pre-expression (green) and Round 0 reinfected post-expression phage (orange).

### Linear epitope profiling by multi-round phage display reveals significant binding to the N-terminal epitope tag

To enrich antigen-specific clones from the antibody repertoire of rabbits immunized with TeeVax3, we first purified a TeeVax3-specific pool of antibodies by immunoprecipitation. High-purity TeeVax3 antigen (Figure 3a) was conjugated to agarose beads to enable the isolation of both linear and conformational TeeVax3 epitope-binding clones. This process yielded approximately 1.3 mg of TeeVax3-specific antibodies per mL of immunized rabbit sera. SDS PAGE confirmed a high purity of the resulting antibody fraction with multiple bands representing heavy and light chains from multiple antibody subclasses (Figure 3b). These antibodies were directly used for phage panning of the TeeVax3 linear epitope library. We performed three rounds of panning and used Nanopore Sequencing to track the clonal enrichment of each tile. Enrichment scores were determined by calculating the ratio of observed tile frequencies from the pre-panned phage library and post-panned libraries.

**Figure 3:**
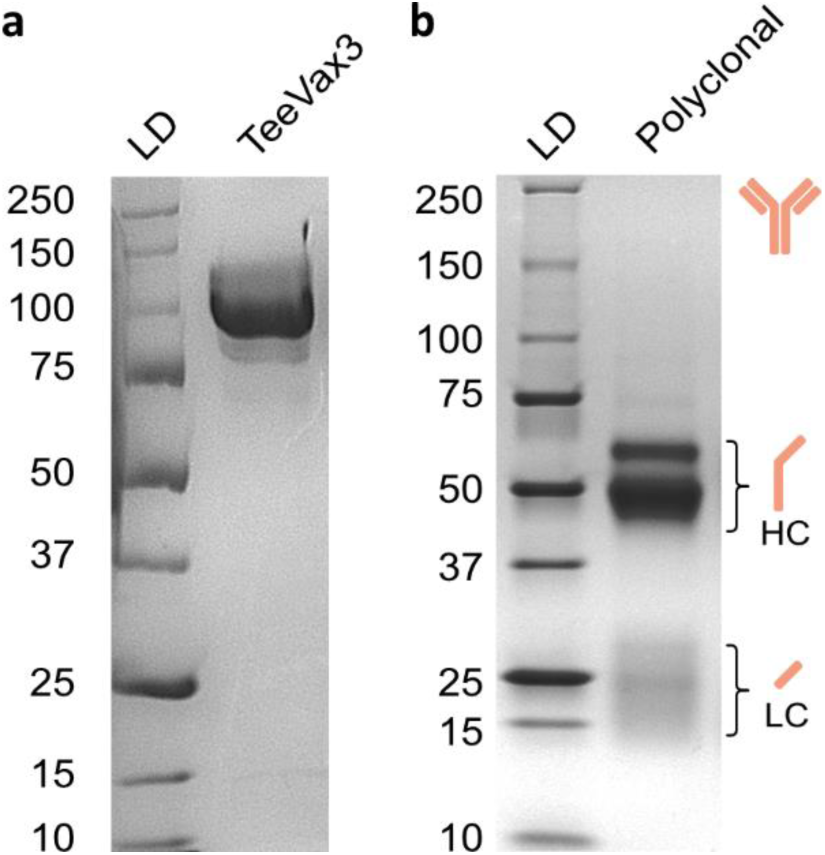
Expression of TeeVax3 antigen and isolation of vaccine-specific antibodies. **(a)** TeeVax3 was expressed in *E. coli,* purified via Ni-NTA affinity chromatography and run on SDS-PAGE. **(b)** A custom chromatographic resin was created by conjugating TeeVax3 antigen to aldehyde-activated agarose beads and used to purify polyclonal antibodies from TeeVax3-immunized rabbits. Separation by reducing SDS-PAGE gel shows the various heavy chain and light chain sizes expected from a diversity of clonal sequences and antibody subclasses.

Over 64,000 high-quality sequences were obtained from phages after each round of panning, representing a greater than 200-fold coverage of the theoretical library diversity. We observed a rapid bias in representation towards tiles from the N-terminus of the TeeVax3 antigen corresponding to the 6-His epitope tag and TEV cleavage sequences (Figure 4a). By round two of panning, the phage library sequence pool was dominated by 81.1% of tile 1 (MSYYHHHHHHDYDIPTTENL) and 9.5% of tile 2 (YHHHHHHDYDIPTTENLYFQ). Little further enrichment was observed in the third round of panning. We next explored the likelihood these sequences were the result of expression bias by comparing the enrichment of these tiles between the Round 0 pre-expression plasmid library and Round 0 reinfected post-expression frequencies. We found that while the proportion of tile 1 increased from 0.347% to 0.645% (1.9-fold), indicating a positive expression bias, tile 2 dropped in abundance from 0.404% to 0.176% (0.44-fold) indicating expression of this sequence was relatively disfavored. In each case, the >10-fold enrichment of these sequences over the first round of panning far exceeds expression bias contributions. More broadly, we observed a decrease in the median tile frequency from 0.328% to 0.260% across all the tiles.

**Figure 4:**
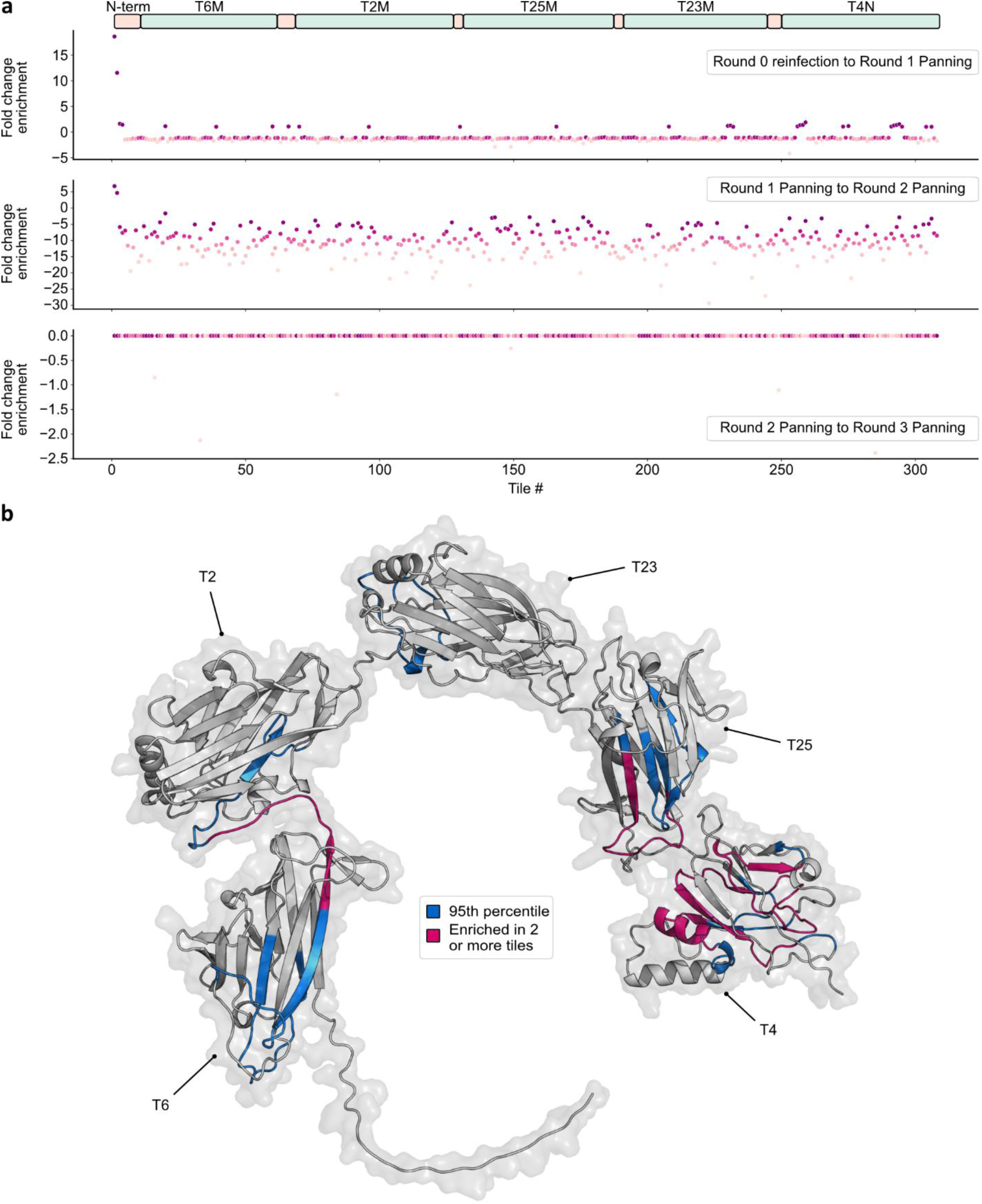
Tile enrichment during panning against TeeVax3-specific antibody pool. **(a)** The TeeVax3 peptide pool was expressed on bacteriophage and panned against immobilized anti-TeeVax3 polyclonal antibodies. Fold enrichment values for each peptide tile were computed by dividing the post-panning frequency by the corresponding pre-panning frequency following Nanopore sequencing. Each data point represents an individual peptide tile sequence. **(b)** After removing the high abundance N-terminal tiles (1-3), tiles above the 95^th^ percentile were mapped to a model of TeeVax3. Blue indicates tiles above the 95^th^ percentile and magenta shows sequences represented in two or more tiles.

To further investigate clonal bias outside of the N-terminal sequences, we expanded the analysis between the Round 0 reinfected library and Round 1 of panning to include tiles with enrichment in the 95^th^ percentile exclusive of tiles 1-3. These enriched peptides were mapped onto a three-dimensional representative model of the TeeVax3 antigen, generated using AlphaFold2^26^ in the absence of crystal structure data (**Supplementary Fig. S2**). This process revealed a lack of antibody binding preference for the T2M domain, T25M domain, and the inter-domain spacer regions, except for the linker between the T6M and T2M domains. Instead, we observed some preference for T6M, T23M, and increasingly T4N binding where a high proportion of overlapping tiles were mapped (Figure 4b).

### High-resolution epitope profiling defines a high-affinity peptide within the epitope tag

There remained the possibility that the unexpected binding we observed for the epitope tag could be a small fraction of the total antibody pool with most of the response being dominated by conformational epitope binders rather than linear epitope binders. To measure the relative abundance of linear binders to confirmation binders, we cleaved the His tag from TeeVax3 using TEV protease and confirmed the absence of the His/TEV tag by ELISA using anti-His HRP for detection (Figure 5a). ELISA analysis was then used to measure the relative anti-TeeVax3 antibody signal from purified sera samples with and without the His/TEV tag. We observed a decrease in antibody EC_50_ (6.3-fold) upon removal of the epitope tag indicating the binders to Tiles 1 and 2 dominated the pool of antibodies against the antigen even with conformational binding clones present (Figure 5b).

**Figure 5:**
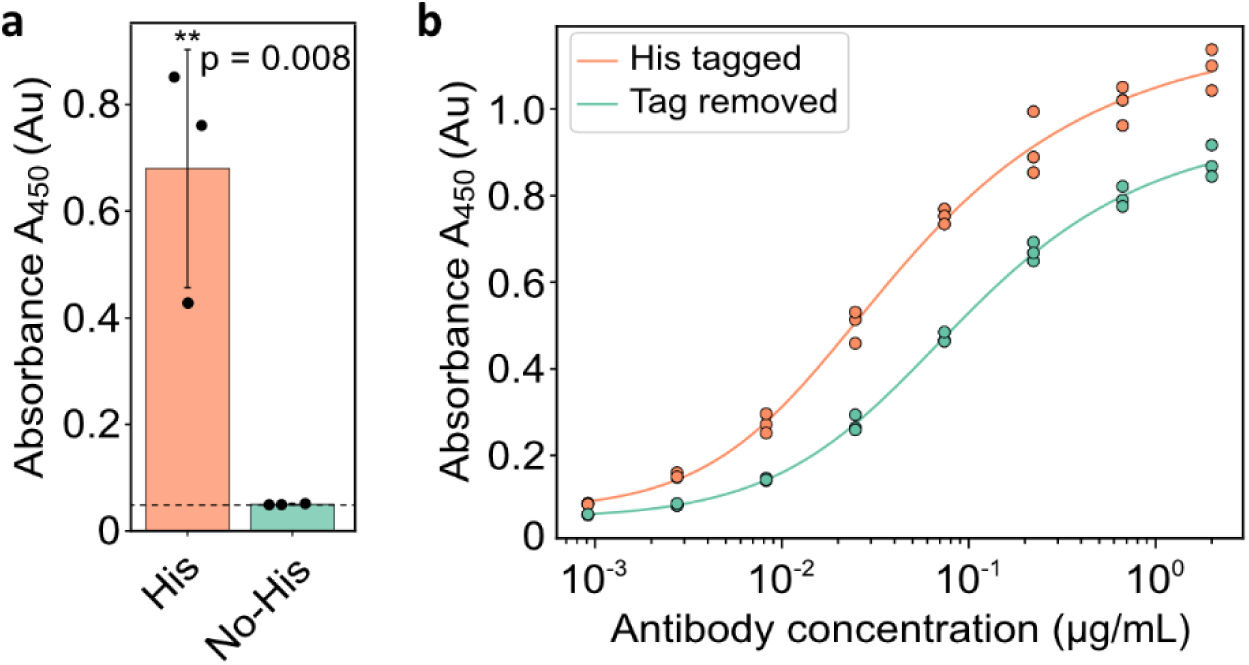
Antibody response with and without the N-terminal epitope tag sequence. **(a)** TeeVax3 antigens were digested with TEV protease and the presence of residual Histidine epitope tag was assayed by single-well ELISA relative to the undigested antigen. The dashed line indicates background binding in the assay. Data are presented as the mean ± SD (n = 3). **(b)** Representative ELISA plot showing polyclonal antibody responses towards TeeVax3 antigen with and without the epitope tag removed (n=3).

**Figure 6:**
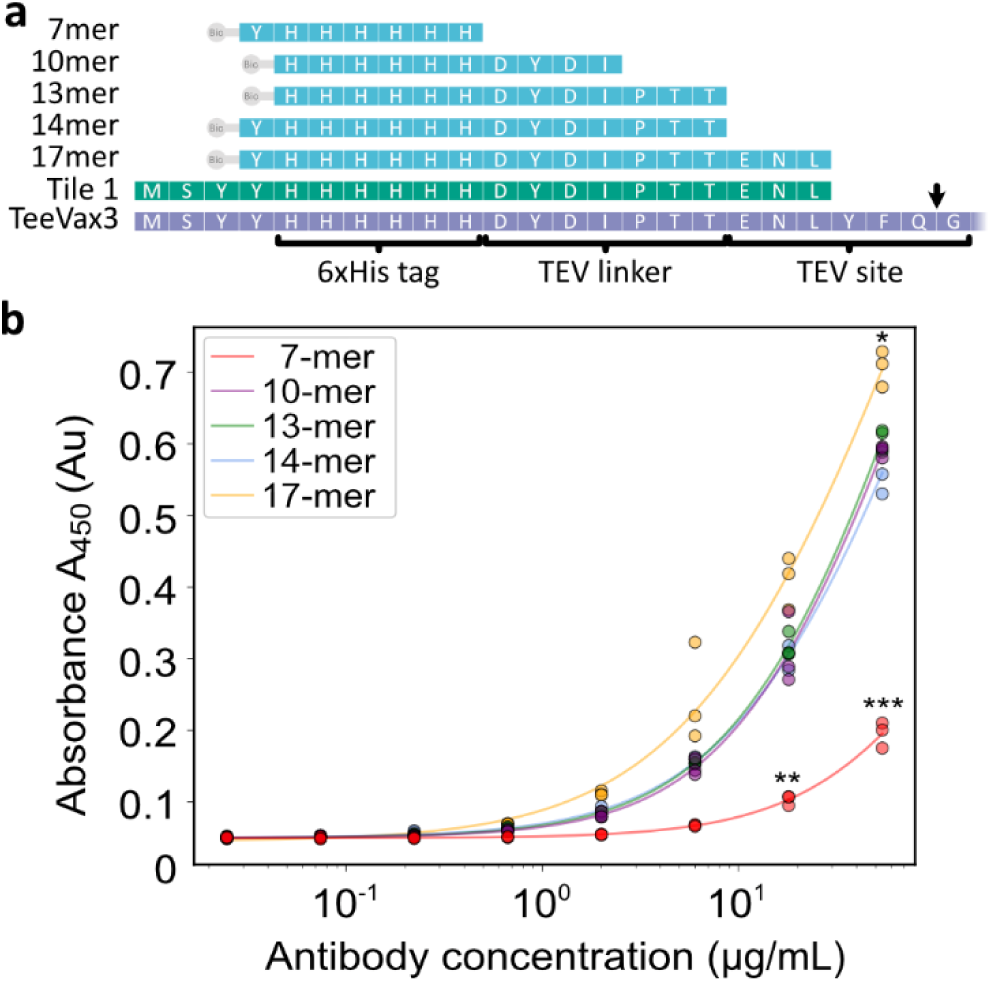
Fine epitope profiling with synthetic peptides. **(a)** Peptide series with N-terminal biotin fusions generated from the Tile 1 TeeVax3 sequence. **(b)** ELISA analysis measuring the binding capacity of polyclonal TeeVax3-specific antibodies against biotinylated peptide series. Pairwise t-tests were used for pairwise comparison at each concentration with Bonferroni correction for multiple comparisons (*** p < 0.001, ** p < 0.01, * p < 0.05).

The unexpected immunogenicity of the epitope tag led us to profile the specific amino acids driving the antibody response with greater precision. We ordered a series of synthetic biotinylated peptides of decreasing size beginning with the 17 amino acids overlapping in Tiles 1 and 2 that provided the greatest enrichment by phage panning. Subsequent peptides reduced the size of this epitope with the smallest fragment tested consisting of a 7 amino acid sequence encompassing the 6-His tag (Figure 5a). The peptides were captured with neutravidin and TeeVax3-specific antibodies were assayed for binding. We found the highest relative signal with the longest 17 amino acid epitope and a trend of similar binding for each of the 14-mer, 13-mer, and 10-mer tiles (Figure 5b). A substantial drop in signal was observed for the shortest tile covering the 6-His tag with an additional Tyrosine motif. This dataset suggests the strongly immunogenic motif we observe encompasses both residues from the Histidine epitope tag and the TEV linker regions.

## Discussion

High-throughput epitope mapping tools provide a rapid snapshot of antibody binding bias against vaccine or pathogen antigens. In this study, we explored the use of Nanopore sequencing to rapidly profile linear epitope antibody binders derived from rabbit sera against the TeeVax3 candidate vaccine for Group A Streptococcus. An oligo pool was ordered and used to create a library of 20-mer peptides derived from the TeeVax3 antigen. We then used a multi-round phage panning approach against enriched TeeVax3-specific polyclonal antibodies to monitor the enrichment of the expressed tiled peptides. After three rounds of panning, the resulting libraries were dominated by peptide tiles stemming from the N-terminus. We were surprised to find a strong enrichment of peptide sequences encoding the 6 His/TEV purification tag and therefore evaluated the relative binding signal of antibodies against the TeeVax3 antigen with and without epitope tag cleavage. This analysis confirmed the polyclonal pool was dominated by linear binders to sequences from the epitope tag and confirmed the unexpected immunogenicity observed by phage panning. To further understand the scope of the immunogenic epitope, we ordered a series of biotinylated peptides of decreasing size and observed binding even for the 6-His tag with one additional residue. Our findings indicate the residues displaying the strongest immunogenicity lie within a 10 amino acid sequence covering the 6-His epitope tag and the N-terminus of the TEV linker sequence.

Prior studies have used sequencing approaches with very low error rates, such as Illumina^27,28^, for linear epitope binding antibody mapping. The accessibility and speed of Nanopore allows for in-laboratory sequencing even if core facilities or next-generation sequencing providers are not readily available. Despite a high error rate, we found the sequence distance generated by offset peptide tiles was sufficient for accurate binning and determination of read counts required to calculate enrichment factors. While we have used only a limited set of peptide tiles here (308) relative to other examples in literature^29,30^, the length of 20 amino acids means it is very unlikely sequencing errors could cause binning errors in larger libraries, with the exception of antigens with repeats or regions of high sequence homology. The finding that most antibody binders were directed to the epitope tag was unexpected, especially considering that His tags are typically considered to have low immunogenicity. While there are reports of His tags influencing antigen immunogenicity^31^, raising antibodies against His tags frequently requires conjugation to carrier proteins to elicit sufficient antibody titers^32^. It should be noted the strongest antibody binding signal we observed did require residues from the TEV cleavage site in addition to those from the 6-His tag. An obvious mitigation strategy is to remove the His tag via TEV cleavage, but this step is inconsistently performed in preclinical vaccine studies. The generation of non-productive antibodies against epitope tags could impact observed vaccine efficacy in early phases. It is also worth mentioning that TEV cleavage leaves a residual serine residue (ENLYFQ\S) that is not likely to be immunogenic by itself but could contribute to the development of a neoepitope.

A further challenge with using multiple rounds of phage panning is the resulting enrichment of high expressing rather than high-affinity peptide clones. Sequences with a growth advantage can bias tile frequency and lead to the false positive identification of linear epitopes. Here, we took steps to mitigate this issue by monitoring tile abundance before and after phage expression as well as using synthetic peptides to validate our top N-terminal hits. We suggest a maximum of two rounds of panning are required to detect tile enrichment when using next-generation sequencing approaches, as strong binders rapidly dominate the library. Further, employing two rounds of panning could provide additional information on the relative affinity of interactions assuming a limited influence from expression bias. When we removed the high abundance epitope tag peptides from the analysis and mapped the remaining tiles above the 95^th^ percentile, we found a small number of linear epitopes unevenly distributed across the TeeVax3 antigen. Previous work has also found biases in antibody binding to T-antigens upon immunization. Raynes et al., report conformational binders with a bias towards the N-terminal ends of T18.1 N-domains exposed in a temperature-dependent manner ^33^. While these data are from a limited subset of T-antigens it is clear further exploration of immunodominant motifs would be valuable for developing these proteins as vaccines.

Epitope mapping studies provide insights into immune response bias that inform antigen design in vaccines. Here, we identify the epitope tag on TeeVax3 as having unanticipated immunogenicity. Beyond the scope of the findings presented here, our method could be used to further profile larger peptide libraries. For example, to establish TeeVax3 correlates of protection across the 21 known T antigen groups encoded by the *tee* gene from Group A Strep pathogens^34,35^. This data would best be paired with experimentally generated information on conformational epitope binding. Nonetheless, the method presented here provides an accessible and low-cost approach to linear epitope panning using tools available to most molecular biology laboratories.

## Methods

### Antibody and antigen purification

The TeeVax3 antigen was expressed in *E. coli* by adapting previously described methods^15^. Briefly, pROEXhtb vectors encoding TeeVax3 were transformed into BL21 (DE3) cells and cultures in exponential growth phase induced with 0.1 mM Isopropyl β-d-1-thiogalactopyranoside (IPTG). After overnight growth at 18 °C for 18 hours, cell pellets were resuspended in Lysis buffer (50 mM Tris-HCl pH8, 150 mM NaCl, 20 mM imidazole), sonicated, and centrifuged to recover the supernatant. His tagged antigens were purified by Ni-NTA chromatography using 5 mL HisTrap HP columns (Cytiva) on a BioRad NGC FPLC. Non-specifically bound proteins were removed with 50 mM Tris-Cl pH 8, 150 mM NaCl, 20 mM imidazole and the antigen eluted with 50 mM Tris-Cl pH 8, 150 mM NaCl, 1 M imidazole. Amicon Ultra-15 10 kDa centrifugal filters (Merck Millipore) were used to concentrate the protein before size exclusion polishing with a S75 16/60 column (GE Healthcare) using 50 mM Tris (pH 8), 150 mM NaCl. Samples were buffer exchanged into 1x PBS for storage and analysed by 12% SDS-PAGE gel electrophoresis. For His tag removal, TeeVax3 antigen was buffer exchanged into 20 mM Tris pH 7.5 and digested overnight at 4 °C with TEV Protease (NEB). Undigested antigen and digested His tags were captured by Ni-NTA chromatography as described above, taking the flowthrough fraction as the untagged antigen.

TeeVax3 immunized rabbit sera were obtained from New Zealand white (NZW) rabbits immunized subcutaneously with 100 μg of recombinant protein emulsified 1:1 with incomplete Freund’s adjuvant and boosted at 2 and 4 weeks. Antiserum was collected on day 42. All animal experiments were approved by The University of Auckland Animal Ethics Committee and performed in the Vernon Jansen Unit at The University of Auckland. Immunoprecipitation of TeeVax3-specific antibodies was performed by coupling TeeVax3 antigen to AminoLink Plus Coupling Resin (ThermoFisher). 8 mg of TeeVax3 antigen in 0.1 M sodium phosphate, 0.15 M NaCl, pH 7.2 coupling buffer were incubated with 2 mL of resin at 4°C overnight with gentle rotation. The following day, 5 mL of pH 7.2 PBS and 100 µL of Cyanoborohydride Solution were added and the mixture rotated for 4 hr at room temperature. 1 mL of conjugated resin was added to a gravity flow column and sera diluted five-fold in PBS was passed through the column. The bound fraction was washed with PBS, eluted with 0.1 M glycine pH 3, and neutralized with 10% the solution volume of 1M Tris pH 9. Amicon Ultra-15 10 kDa centrifugal filters (Merck Millipore) were used for buffer exchange into PBS prior to purity analysis by SDS PAGE. Purification of Teevax3-specific antibodies was performed by diluting serum five-fold in PBS and passing it through the conjugated resin. The bound antibody pool was washed with at least 10 CV of PBS and eluted with 0.1 M glycine pH 3. The pH was raised with 1/10^th^ the volume of 1M Tris pH 9, buffer exchanged into PBS and analyzed by 12% SDS PAGE.

### Antigen tiling

The tile library was created computationally by splitting the 941 amino acid sequence of the Teevax3 antigen (**Supplementary Table S1**) into 308 segments each 20 amino acids long. These peptides were offset by three amino acids to ensure complete coverage of potential binding sites. Bioinformatics software (Geneious Prime) was then used to back translate the tiles into nucleotide sequences, followed by the addition of adaptors containing a coding sequence, peptide transport signal and restriction sites to both 5’ and 3’ ends (5’ adaptor CGAGGGCCCAGCCGGCCATGGCCGAGGGT, 3’ adaptor TCGGCCTCGGGGGCCA). The resulting 308 tiles of 105 nucleotides each were then ordered as a single oligo pool (Twist Bioscience).

### Phage library construction and expression

The oligo pool was duplexed by a single cycle of PCR using OneTaq® Quick-Load® 2X Master Mix with Standard Buffer (NEB). A reverse primer (5’-TGGCCCCCGAGGCC-3’) was included in the reaction and the following conditions were used: denaturation at 94 °C for 1 min, 1 cycle of elongation at 60 °C for 1 min, followed by final extension for 5 min at 68 °C. The double-stranded oligo pool was digested with Sfil restriction enzyme for 2 hours at 50 °C. Digested fragments were cleaned using a QIAquick nucleotide removal kit (Qiagen) and DNA concentration was quantified using a DeNovix double-stranded Ultra sensitivity assay. The library was then ligated into SfiI digested pAK200 vector and bound to Monarch columns (NEB) after the addition of five volumes of 5 M Guanidine Hydrochloride containing 30% isopropanol. The sample was then washed with 10 mM Tris-HCl pH 7.5, 80% ethanol and eluted with ultra-pure water. Ligated plasmids were transformed into electrocompetent SS320 cells and plated to create the final library. Library glycerol stocks were grown at 37 °C from OD 0.1 to 0.4-0.6 in LB media supplemented with 20 ug/mL chloramphenicol. M13K07 helper phage were added at an MOI of 20x and the culture incubated without shaking for 1 hour to allow infection. Kanamycin antibiotic was added (35 ug/mL) and peptide expression was induced with 1 mM IPTG before overnight growth to produce phage. Culture supernatants were filtered through 0.22 µm filters and phage were precipitated by adding 20% of the total volume of 20% PEG/2.5 M NaCl and incubating on ice for at least 30 min. Phage were centrifuged at 16000 g, resuspended in 1x PBS and spun again twice more at 16,000 g changing the tube between spins. Titers were quantified by spectrophotometry (DeNovix DS11) using the formula *virions/mL* = (A_269_ – A_320_) × 6 × 10^16^/(*phage ssDNA bases*)^36^.

### Phage panning

Phage panning was carried out in ELISA 8 well strips (Corning). The wells were coated overnight at 4°C with 100 µL of purified anti-TeeVax3 antibodies at 4 µg/mL in 50 mM Na_2_CO_3_ buffer at pH 9.5 or incubated with Na_2_CO_3_ buffer alone. The next day, the solutions were removed, and the plates were blocked with 300 µL PBSMT (PBS containing 2% skim milk powder and 0.05% Tween20). After 1 hour of shaking at room temperature, the strips were washed 3x with PBST (PBS containing 0.05% Tween20). 1 x 10^11^ phage per well were then added for 1 hour to the strips coated with buffer alone for non-specific phage binding depletion before transfer to the wells coated with antigen. The wells were washed again after 1 hour, although the number of washes was increased by three each round to increase selection pressure. Phages were eluted with a 10 min incubation using 100 µL of 0.1 M glycine/HCl pH 2.7, transferred to a 2 mL tube previously blocked with PBS containing 2% skim milk powder, and neutralized with 1vM Tris pH 8.0 at a 1:5 ratio (vol Tris/vol phage). 20 mL SS320 E. coli cells were grown in 2xYT media and 2% glucose medium to an OD 0.4-0.6 at 37°C without shaking, and incubated with the eluted phages for 30 min at 30 °C. The shaking speed was then increased to 250 rpm for another 30 min. The phage library was recovered by plating infected cells on three 145 mm round LB agar plates containing 2% glucose and 20 µg/mL chloramphenicol and overnight growth at 37°C.

### Nanopore sequencing and analysis

Library plasmid stocks from each screening phase were directly purified from corresponding *E. coli* glycerol stocks using via QIAprep Spin Miniprep Kit (Qiagen, Germany). Linear, 723 bp DNA fragments representing the tile library CDS were gel excised and purified with a QIAquick Gel Extraction Kit (Qiagen) following restriction digest of each library plasmid stock with *Bam*HI-HF and *Xba*I (NEB, USA). Each purified library fragment was then prepared according to the SQK-NBD114.24 Oxford Nanopore Technologies (ONT, UK) ligation sequencing amplicons protocol and kit (200 fmol input), and subsequently sequenced using a R10.4.1 flow cell (ONT). Sequencing was stopped once the estimated raw read count reached ≥100x coverage per tile per sample. Base-calling and demultiplexing of raw read data was performed using Dorado (v0.5.3; model dna_r10.4.1_e8.2_400bps_sup@4.3.0). Read data were filtered for quality control using fastp (v0.23.4^37^) and binned into tile counts via local alignment against the 308 segments using minimap2 (v2.24^38^) and samtools (v1.19.2^39^).

### ELISA analysis

ELISA analysis was used to compare antibody binding to TeeVax3 with and without the His epitope tag. Antigens were coated overnight at 4 °C on ELISA plates (Corning 3590) at 4 µg/ml in Coating Buffer (50 mM NaHCO3, pH 9.5), washed the next day three times with PBS containing 0.05% Tween20, and blocked with PBS containing 2% w/v skim milk powder. Blocked plates were washed a further three times with PBS Tween20. Serial dilutions of TeeVax3-specific antibodies isolated from rabbit sera were added to the plate, and a goat anti-rabbit IgG HRP antibody (Abcam) was used at 1:50,000 for detection of binding. Control samples confirming His tag removal from the TeeVax3 antigen used a HRP conjugated anti-polyHistidine antibody [HIS-1, Abcam] diluted 1:20,000 in PBSM. 50 µl 1-Step™ Ultra TMB-ELISA (ThermoFisher) was used for detection of HRP signal in each well and the reaction quenched after 10-15 min with an equal volume of 1 M H_2_SO_4_ before reading at 450 nm on a Spectramax M4. All ELISA steps were undertaken at room temperature with incubation periods of 1 hr unless otherwise indicated. Fine epitope mapping was performed by ordering synthetic peptides with N-terminal biotin residues (Genscript: 17-mer YHHHHHHDYDIPTTENL, 14-mer YHHHHHHDYDIPTT, 13-mer HHHHHHDYDIPT, 10-mer HHHHHHDYDI, 7-mer YHHHHHH). ELISA analysis followed the protocol above except the ELISA plates were coated with biotinylated peptides that had been precomplexed with neutravidin at a 4:1 molar ratio for 30 min at room temperature in 50 mM NaHCO3, pH 9.5. ELISA data was fit by non-linear regression of a reparametrized 5 parameter logistic model in JupyterLab^40^.

## Acknowledgements

This work was supported by an MWC project grant (#4026) that also supported K.BC with a Masters Scholarship. K.H. was supported by a University of Waikato Doctoral scholarship. W.K. was supported by a New Zealand Marsden Grant (23-UOW-006).

## Author contributions

K.BC. and K.H. contributed equally to this work. K.H., J.L., and W.K. conceived of and designed the study. K.BC., K.H., K.B., and M.R. performed laboratory experiments and created reagents used in the study. K.H. and K.BC. performed the Nanopore sequencing and bioinformatic analysis. K.H, K.BC., and W.K. wrote the manuscript. All authors provided feedback and revisions for the final version of the manuscript.

## Data availability statement (mandatory)

Processed data and scripts used in this work can be found at https://github.com/ImmunoLab/LinPepEpitopes. Raw Nanopore sequencing data is available in the NCBI SRA database under BioProject 1195632 (http://www.ncbi.nlm.nih.gov/bioproject/1195632).

## Additional Information (including a Competing Interests Statement)

The authors declare no competing interests.

## References

1. Larocca, R. A. et al. Vaccine protection against Zika virus from Brazil. Nature 536, 474–478 (2016).

2. Liu Yihao et al. Robust induction of B cell and T cell responses by a third dose of inactivated SARS-CoV-2 vaccine. Cell Discov. 8, 10 (2022).

3. McNamara, H. A. et al. Antibody Feedback Limits the Expansion of B Cell Responses to Malaria Vaccination but Drives Diversification of the Humoral Response. Cell Host Microbe 28, 572–585.e7 (2020).

4. Cao, Y. et al. BA.2.12.1, BA.4 and BA.5 escape antibodies elicited by Omicron infection. Nature 608, 593–602 (2022).

5. Sevvana, M. & Kuhn, R. J. Mapping the diverse structural landscape of the flavivirus antibody repertoire. Curr. Opin. Virol. 45, 51–64 (2020).

6. Cia, G., Pucci, F. & Rooman, M. Critical review of conformational B-cell epitope prediction methods. Brief. Bioinform. 24, bbac567 (2023).

7. Hu, D. & Irving, A. T. Massively-multiplexed epitope mapping techniques for viral antigen discovery. Front. Immunol. 14, 1192385 (2023).

8. De Leon, A. J., Tjiam, M. C. & Yu, Y. B cell epitope mapping: The journey to better vaccines and therapeutic antibodies. Biochim. Biophys. Acta BBA - Gen. Subj. 1868, 130674 (2024).

9. Mohan, D. et al. PhIP-Seq characterization of serum antibodies using oligonucleotide-encoded peptidomes. Nat. Protoc. 13, 1958–1978 (2018).

10. Antanasijevic, A. et al. From structure to sequence: Antibody discovery using cryoEM. Sci. Adv. 8, eabk2039 (2022).

11. Shrock, E. L., Shrock, C. L. & Elledge, S. J. VirScan: high-throughput profiling of antiviral antibody epitopes. Bio-Protoc. 12, e4464–e4464 (2022).

12. Xu, G. J. et al. Viral immunology. Comprehensive serological profiling of human populations using a synthetic human virome. Science 348, aaa0698 (2015).

13. Garrett, M. E. et al. Phage-DMS: A Comprehensive Method for Fine Mapping of Antibody Epitopes. iScience 23, 101622 (2020).

14. Hanning, K. R., Minot, M., Warrender, A. K., Kelton, W. & Reddy, S. T. Deep mutational scanning for therapeutic antibody engineering. Trends Pharmacol. Sci. 43, 123–135 (2022).

15. Loh, J. M. et al. A multivalent T-antigen-based vaccine for Group A Streptococcus. Sci. Rep. 11, 4353 (2021).

16. Fan, J., Toth, I. & Stephenson, R. J. Recent Scientific Advancements towards a Vaccine against Group A Streptococcus. Vaccines 12, 272 (2024).

17. Raynes, J. M. et al. Identification of an immunodominant region on a group A Streptococcus T-antigen reveals temperature-dependent motion in pili. Virulence 14, 2180228 (2023).

18. López-Laguna, H. et al. Insights on the emerging biotechnology of histidine-rich peptides. Biotechnol. Adv. 54, 107817 (2022).

19. Zhao, X., Li, G. & Liang, S. Several Affinity Tags Commonly Used in Chromatographic Purification. J. Anal. Methods Chem. 2013, 1–8 (2013).

20. Rodrigues, M. X., Yang, Y., de Souza Meira, E. B., do Carmo Silva, J. & Bicalho, R. C. Development and evaluation of a new recombinant protein vaccine (YidR) against Klebsiella pneumoniae infection. Vaccine 38, 4640–4648 (2020).

21. Otsyula, N. et al. Results from tandem Phase 1 studies evaluating the safety, reactogenicity and immunogenicity of the vaccine candidate antigen Plasmodium falciparum FVO merozoite surface protein-1 (MSP142) administered intramuscularly with adjuvant system AS01. Malar. J. 12, 29 (2013).

22. Gu, H. et al. Adaptation of SARS-CoV-2 in BALB/c mice for testing vaccine efficacy. Science 369, 1603– 1607 (2020).

23. Ngampasutadol, J., Rice, P. A., Walsh, M. T. & Gulati, S. Characterization of a peptide vaccine candidate mimicking an oligosaccharide epitope of *Neisseria gonorrhoeae* and resultant immune responses and function. Vaccine 24, 157–170 (2006).

24. Krebber, A. et al. Reliable cloning of functional antibody variable domains from hybridomas and spleen cell repertoires employing a reengineered phage display system. J. Immunol. Methods 201, 35–55 (1997).

25. Gulati, S., McQuillen, D. P., Mandrell, R. E., Jani, D. B. & Rice, P. A. Immunogenicity of Neisseria gonorrhoeae Lipooligosaccharide Epitope 2C7, Widely Expressed In Vivo with No Immunochemical Similarity to Human Glycosphingolipids. J. Infect. Dis. 174, 1223–1237 (1996).

26. Jumper, J. et al. Highly accurate protein structure prediction with AlphaFold. nature 596, 583–589 (2021).

27. Qi, H. et al. Antibody Binding Epitope Mapping (AbMap) of Hundred Antibodies in a Single Run. Mol. Cell. Proteomics 20, 100059 (2021).

28. Paull, M. L., Bozekowski, J. D. & Daugherty, P. S. Mapping antibody binding using multiplexed epitope substitution analysis. J. Immunol. Methods 499, 113178 (2021).

29. Rajan, J. V. et al. Phage display demonstrates durable differences in serological profile by route of inoculation in primary infections of non-human primates with Dengue Virus 1. Sci. Rep. 11, 10823 (2021).

30. Yaffe, Z. A. et al. Passively Acquired Constant Region 5–Specific Antibodies Associated With Improved Survival in Infants Who Acquire Human Immunodeficiency Virus. Open Forum Infect. Dis. 10, ofad316 (2023).

31. Singh, M. et al. Effect of N-terminal poly histidine-tag on immunogenicity of *Streptococcus pneumoniae* surface protein SP0845. Int. J. Biol. Macromol. 163, 1240–1248 (2020).

32. Przedpelski, A., Tepp, W. H., Pellett, S., Johnson, E. A. & Barbieri, J. T. A Novel High-Potency Tetanus Vaccine. mBio 11, e01668–20 (2020).

33. Raynes, J. M. et al. Identification of an immunodominant region on a group A Streptococcus T-antigen reveals temperature-dependent motion in pili. Virulence 14, 2180228 (2023).

34. Steemson, J. D. et al. Survey of the bp/tee genes from clinical group A streptococcus isolates in New Zealand – implications for vaccine development. J. Med. Microbiol. 63, 1670–1678 (2014).

35. Falugi, F. et al. Sequence Variation in Group A Streptococcus Pili and Association of Pilus Backbone Types with Lancefield T Serotypes. J. Infect. Dis. 198, 1834–1841 (2008).

36. Smith, G. P. & Scott, J. K. Libraries of peptides and proteins displayed on filamentous phage. In Methods in Enzymology vol. 217 228–257 (Academic Press, 1993).

37. Chen, S. Ultrafast one-pass FASTQ data preprocessing, quality control, and deduplication using fastp. iMeta 2, e107 (2023).

38. Li, H. New strategies to improve minimap2 alignment accuracy. Bioinformatics 37, 4572–4574 (2021).

39. Danecek, P. et al. Twelve years of SAMtools and BCFtools. GigaScience 10, giab008 (2021).

40. Liao, J. J. Z. & Liu, R. Re-parameterization of five-parameter logistic function. J. Chemom. 23, 248–253 (2009).

