## Supplemental material for "A rapid approach for linear epitope vaccine profiling reveals unexpected epitope tag immunogenicity"

**Supplementary Material**

### Supplementary Tables

**Table S1. Amino acid sequence of the TeeVax3 antigen.** The underlined sequence is the N-terminal epitope tag

| | |
|---|---|
| >TeeVax3 | MSYYHHHHHHDYDIPTTENLYFQGAMGSPSITKKVTGTIDDVNKKTTSLGSVLSYSLT<br>FELPSYTKEAVNKTVYVSDNMSEGLTFNFNSLTVEWKGKMANITEDGSVMVENTKIG<br>IAKEVNNGFNLSFIYDSLESISPNISYKAVVNNKAIVGEEGNPNKAEFFYSNNPTKGNTY<br>DNLDKKPDKGNGITSKEDSKIVYTYRSGSGSGKNTTVNEPKVDKDVTKLGKDDDTYQ<br>IGDKITWFLKSTVPSNIKTLDKFGFTDTLNKGLSFIGDKTQTVTKVQFGTTVLSPDTDY<br>TVEILDSKLTVSLTSAGIEKVSGLVASKQLITEAEKLYKAEDNTDEAAFLSVEVNAKLN<br>ADAVMGSRIENDVELDYGHESDIYKSKVPTNEVPEVHTRSGSGSGPKPGKDVKELGL<br>NHSSYNIGERFSWFLKGTVPKNMLDYEKYSFTDTLDSQLDFISVKSVKYGSQILEKNN<br>DYTLSEPTAQNRTLKVELTEAGIKKVAGLYPDRQEVLDTEIEAIKENTDQKPFLEVEFE<br>TNINSTVILGKPVTNEVKIEFDNKPDKIAKPVTTPPSDNPEVHTRSGSGSGPTISKSITKS<br>TKDGDKDTASVGEKVDYKLTVQLPSYSKDAINKTVFITDKLSQGLTFLPKSLKIIWNG<br>QTLTKVNEEFKAGDKVIAQLKVENNGFNLNFNYDNLDNHAPEVNYSALLNENAVVG<br>KGGNDNNVDYYYSNNPNKGETHKTTEKPKEGEGTGITKKTDKKTVYTYRSGSGSGES<br>SHKTDVVIHKIKMTSLKGWPKEKNPDGTYTGLGDKNYNGEKIDTITSYFGEGAEELDG<br>VSFTYWSVDKEKYKKLTKNPQNYDTVPKMKAFLQGTEKNKALENSSETIDGKTTGHT<br>ADKGGVKVKDLADGYYWFVENSGSNIANGETLSSSAAVPFGLELPVYKADGSTITEL<br>HVYPKNTTTKRS* |

### Supplementary Figures

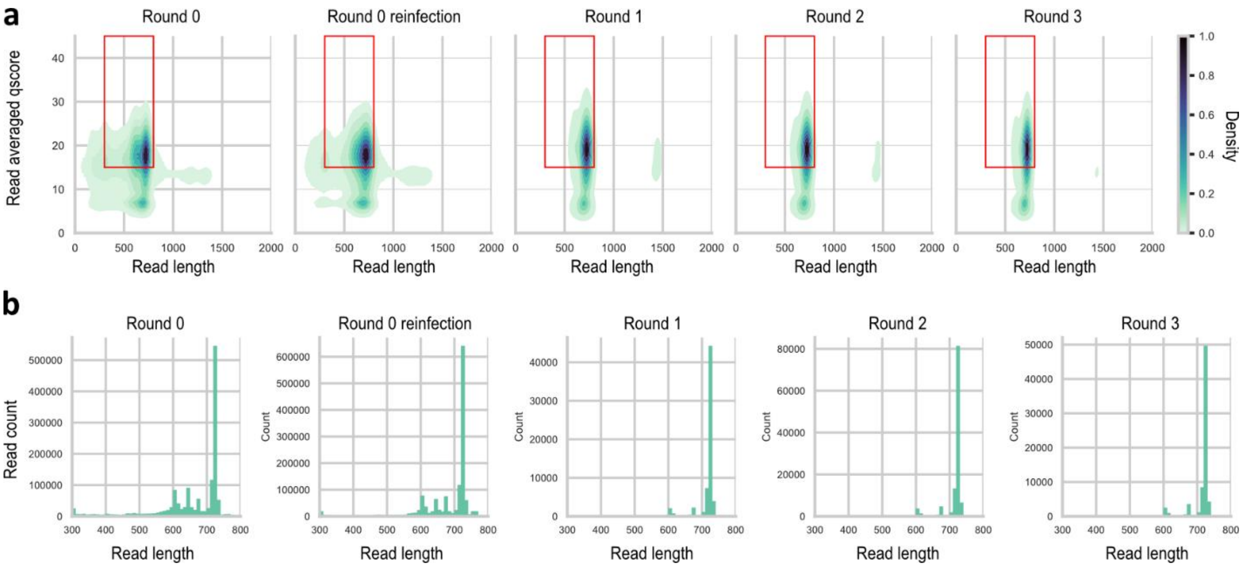


**Figure S1. Nanopore sequencing QC data. (a)** Density plots of qscores averaged across each read are plotted against read lengths for pre-panning libraries and for each round of panning. The red box indicates the data selected for analysis. (b) Histograms of read counts at each read length for pre-panning libraries and for each round of panning.

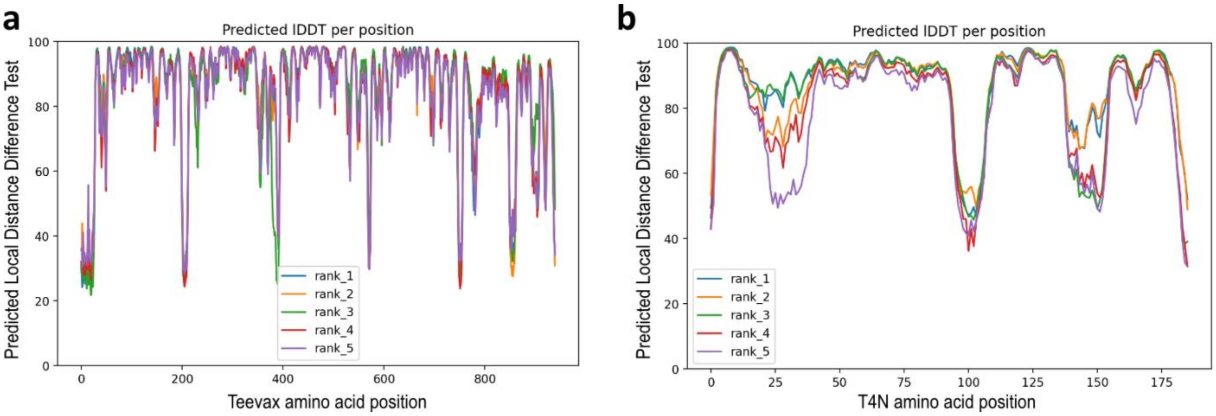


**Figure S2. Structural models of TeeVax3 produced using AlphaFold2 software. (a)** AlphaFold prediction confidence (Local Distance Difference Test scores) across the full length TeeVax3 amino acid sequence. **(b)** AlphaFold prediction confidence (Local Distance Difference Test scores) across the T4N domain alone. To improve the accuracy of the C-terminal T4N domain, the PDB structure 3RPK[1] from the Full-Length Major Pilin RrgB from *Streptococcus pneumoniae* was provided as a template and modeled independently. After alignment in Pymol, the rank_1 prediction was used to create a composite model with the other TeeVax3 domains.

### References

1. Paterson, N. G. & Baker, E. N. Structure of the Full-Length Major Pilin from Streptococcus pneumoniae: Implications for Isopeptide Bond Formation in Gram-Positive Bacterial Pili. *PLOS ONE* **6**, e22095 (2011).
